# Capturing effects of blood flow on the transplanted decellularized nephron with intravital microscopy

**DOI:** 10.1101/2021.02.10.430561

**Authors:** Peter R. Corridon

**Affiliations:** Department of Immunology and Physiology, College of Medicine and Health Sciences, Khalifa University of Science and Technology, PO Box 127788, Abu Dhabi, UAE; Biomedical Engineering, Healthcare Engineering Innovation Center, Khalifa University of Science and Technology, PO Box 127788, Abu Dhabi, UAE; Center for Biotechnology, Khalifa University of Science and Technology, PO Box 127788, Abu Dhabi, UAE; Wake Forest Institute for Regenerative Medicine, Medical Center Boulevard, Winston-Salem, NC 27157-1083, USA

**Keywords:** Intravital microscopy, end-stage renal disease, scaffold, decellularization, kidney bioengineering, autologous and allogeneic transplantation

## Abstract

Organ decellularization creates cell-free, collagen-based extracellular matrices that can be used as scaffolds for tissue engineering applications. This technique has recently gained much attention, yet adequate scaffold repopulation and implantation remain a challenge. Specifically, there still needs to be a greater understanding of scaffold responses post-transplantation and ways we can improve scaffold durability to withstand the in vivo environment. Recent studies have outlined vascular events that limit organ decellularization/recellularization scaffold viability for long-term transplantation. However, these insights have relied on in vitro/in vivo approaches that need enhanced spatial and temporal resolutions to investigate such issues at the microvascular level. This study uses intravital microscopy to gain instant feedback on their structure, function, and deformation dynamics. Thus, the objective of this study was to capture the effects of in vivo blood flow on the decellularized glomerulus, peritubular capillaries, and tubules after autologous and allogeneic orthotopic transplantation into rats. Large molecular weight dextran molecules label the vasculature. They revealed substantial degrees of translocation from glomerular and peritubular capillary tracks to the decellularized tubular epithelium and lumen as early as 12 hours after transplantation, providing real-time evidence of the increases in microvascular permeability. Macromolecular extravasation persisted for a week, during which the decellularized microarchitecture was significantly and comparably compromised and thrombosed in both autologous and allogeneic approaches. These results indicate that in vivo multiphoton microscopy is a powerful approach for studying scaffold viability and identifying ways to promote scaffold longevity and vasculogenesis in bioartificial organs.

## 1 Introduction

Despite the global shortage of organs, transplantation is the best option for those with end-stage renal disease (ESRD), owing to the limitations of dialysis[1–3]. The exponential rise in ESRD, characterized by a debilitating and permanent loss of renal function, has heightened the desire for alternatives such as the bioartificial kidney[4, 5]. Whole organ decellularization has been described as one of the most promising ways to construct a bioartificial kidney[6]. Decellularization focuses on extracting the extracellular matrix (ECM) from the native tissue and organ with as many structural and functional clues as possible[7–10]. The ECM can then be employed as a natural template for regeneration, as observed traditionally in commercial substitutes[11–16].

Preserving the vascular architecture is essential as it provides the ideal template for cellular repopulation[17]. Current studies have outlined vascular events that limit whole organ decellularization and recellularization techniques for long-term transplantation. Several microscopic and macroscopic approaches have examined the whole kidney scaffold structure and function in various settings[18–26]. Such investigations have proved that the decellularization process successfully removes remnant native cellular/tissue components and decellularizing agents while maintaining the vascular and ECM architectures in various large and small animal models.

Across the past four decades since this technology was first outlined, many decellularization and recellularization techniques have been proposed, but critical challenges remain[27]. Specifically, it is difficult to remove all cellular components within a whole organ; thus, decellularization thresholds have been proposed to define critical limits for decellularization[28]. Conversely, adequate repopulation of the epithelial compartment still needs to be attainable[29], particularly in complex organs like the kidney. Current recellularization methods are also plagued by events that lead to incomplete reendothelialization, resulting in structural and functional vulnerabilities. The combination of micro-and macro-vascular complications observed with current related renal transplantation regimens is far more likely to occur in scaffolds with altered significant structural integrities derived from decellularization and recellularization. Most models used to examine these structures within the transplantation environment have primarily utilized in vitro/in vivo imaging techniques to investigate endothelial and epithelial layer porosity alterations, which are known to drive scaffold degradation. However, these methods cannot provide adequate spatial and temporal resolutions, and such limitations outline the need for alternative ways to explore these events. Thus, a better understanding of scaffold integrity and functionality is needed to help improve bioartificial organ development.

Advances in fluorescence tagging and modeling methods enable the tracking of specific cellular and vascular compartments. They can be an essential tool for studying different stages of tissue regeneration [26, 30–37]. One way to achieve this may be to use intravital microscopy (IVM), which provides real-time imaging of cellular-level events and a comprehensive picture of living processes to offer several advantages over in vitro, ex vivo, and 3D models[32]. This technique has been applied extensively to investigate in vivo renal dynamics in normal and pathological settings[38–55]. Since IVM also permits deep tissue imaging at a high resolution, it can help instantly gauge the compatibility of different tissue-engineered constructs after the implantation of biomaterials and track their performance as we strive to create long-term transplantable units.

Recently, de Haan et al.[29] have argued that since adequate repopulation of the epithelial compartment remains unattainable, it seems unlikely that recellularized whole kidneys will be the solution to reduce donor organ shortages. This limitation has posed interest in revising decellularization/recellularization strategies and identifying ways to support this process. Based on these challenges and the benefits of IWM, and as a first step, this study was designed to examine further how in vivo blood flow affects the decellularized microarchitecture after autologous and allogeneic orthotopic transplantation, using well-known vascular and cellular probes, and complements our recent work [7]. Overall, this advanced imaging technique may provide novel insight into scaffold viability and identify ways to promote scaffold longevity and vasculogenesis in whole decellularized organs and help reduce donor organ shortages within the foreseeable future.

## 2 Materials and Methods

### 2.1. Experimental Animals

Experiments were performed on male Sprague-Dawley rats that ranged in weight from 200-400 g (Envigo, Indianapolis, IN, USA). The animals were separated into the following groups (n = 4 for each group): rats in group 1 were used to supply native and decellularized kidneys, respectively, for DNA and SDS quantification assays; rats in group 2 were used for intravital microscopic studies of normal kidneys; and rats in groups 3, 4, 5, and 6 were used for intravital microscopic studies of transplanted acellular kidneys at the 0-hour, 12-hour, 24-hour, and 168-hour (1-week) marks respectively. All animals were given free access to standard rat chow and water throughout the study.

### 2.2 Left Radical Nephrectomy

The left kidney was extracted by first anesthetizing an animal with inhaled isoflurane (Webster Veterinary Supply, Devens, MA, USA), 5% oxygen, and then administering an injection of 50 mg/kg of pentobarbital. Each rat was then placed on a heating pad to maintain a core body temperature of roughly 37 °C. Once the animal was fully sedated, its abdomen was shaved and sanitized with Betadine Surgical Scrub (Purdue Products, Stamford, CT, USA). A midline incision was then made to expose and isolate the left kidney. The associated renal artery, renal vein, and ureter were ligated using 4-0 silk (Fine Science Tools, Foster City, CA, USA) to excise the kidney, with intact segments of the renal artery, vein, and ureter, from each rat. The incision was closed, and all animals recovered for roughly two weeks.

### 2.3 Rat Kidney Decellularization and Sterilization

The arteries of the extracted kidneys were cannulated using PE-50 polyethylene catheter tubing (Clay Adams, Division of Becton Dickson, Parsippany, NJ, USA) and a 14-gage cannula and then secured with a 4-0 silk suture. Kidneys were rapidly perfused with 0.5-1 ml of heparinized PBS. After heparinization, the extracted organs were suspended in PBS, and the cannulated renal artery was perfused with a peristaltic pump (Cole-Palmer, Vernon Hills, IL, USA). The kidneys were then perfused via the renal artery at a 4 ml/min rate with 0.5% sodium dodecyl sulfate (Sigma-Aldrich, St. Louis, MO, USA) for a minimal perfusion time of 6 hours, followed by phosphate-buffered saline (PBS) for 24 hours. The scaffolds were then sterilized with 10.0 Ky gamma irradiation.

### 2.4 DNA Quantification

DNA content in native/control kidneys (n = 4) and decellularized kidneys (n = 4) were measured using a Qiagen DNeasy Kit (Qiagen, Valencia, CA, USA). The tissues were initially minced and stored overnight at −80°C. The next day, the tissues were lyophilized to estimate the ratios of ng DNA per mg dry tissue in each group using the Quant-iT PicoGreen dsDNA assay kit (Invitrogen, Carlsbad, CA, USA) and a SpectraMax M Series Multi-Mode Microplate Reader (Molecular Devices, Sunnyvale, CA, USA) to quantify the DNA content within the extracts.

### 2.5 SDS Quantification

Decellularized scaffolds (n = 4) were first minced and then homogenized using a FastPrep 24 Tissue Homogenizer (MP Biochemicals, Santa Ana, CA, USA). The suspension was then digested with proteinase K (Omega Bio-tek, Atlanta, GA, USA) for roughly 1 h at 56 °C. After which 1 ml methylene blue solution (methylene blue 0.25 g/l, anhydrous sodium sulfate 50 g/l, concentrated sulfuric acid 10 ml/l) was added. The samples were then extracted using chloroform, and absorbance measurements were conducted at a wavelength of 650 nm using the microplate reader to quantify the SDS contents within the extracts.

### 2.6 Florescence Nuclear and Vascular Markers

Hoechst 33342 (Invitrogen, Carlsbad, CA, USA) and 150-kDa fluorescein isothiocyanate (FITC)-dextrans (TdB Consultancy, Uppsala, Sweden) were used to visualize nuclei and vascular compartments, respectively, in live rats. The venous injectates were prepared by diluting 50 μl of a 20 mg/ml stock FITC-dextran solution or 30-50 μl of Hoechst in 0.5 ml of saline.

### 2.7 Autologous and Allogeneic Orthotropic Transplantation of Decellularized Rat Kidneys

After sedation, the torso of each rat was shaved and sanitized with Betadine Surgical Scrub (Purdue Products L.P., Stamford, CT, USA). The tail vein of a sedated rat was either moistened with a warmed sheet of gauze or placed into a warm bath. A 25-gauge butterfly needle was inserted into the dilated tail vein and attached to a syringe containing injectates. A bolus of 0.5 ml heparinized saline was infused into the animal, and incisions were then created to expose each recipient’s previously ligated renal artery and vein. After that, the renal artery and vein of the decellularized kidney were anastomosed end-to-end to the remains of the recipient’s respective clamped left renal artery vein using micro serrefine vascular clamps and microsurgical needles attached to10-0 silk suture thread (Fine Science Tools, Foster City, CA, USA), as presented in the illustration of the harvesting, decellularization, and transplantation process (Figure 1). The ureter was also connected to a proximal portion of the renal vein using the 10-0 silk suture to maintain blood flow patterns across the organ. Other boluses of 0.25-0.5 ml of Hoechst 33342 and 150-kDa FITC-dextran molecules were introduced systemically, and then vascular clamps were removed to establish blood flow into the acellular organ. The transplanted organs were exteriorized for imaging at various measurement time points.

**Figure 1.**
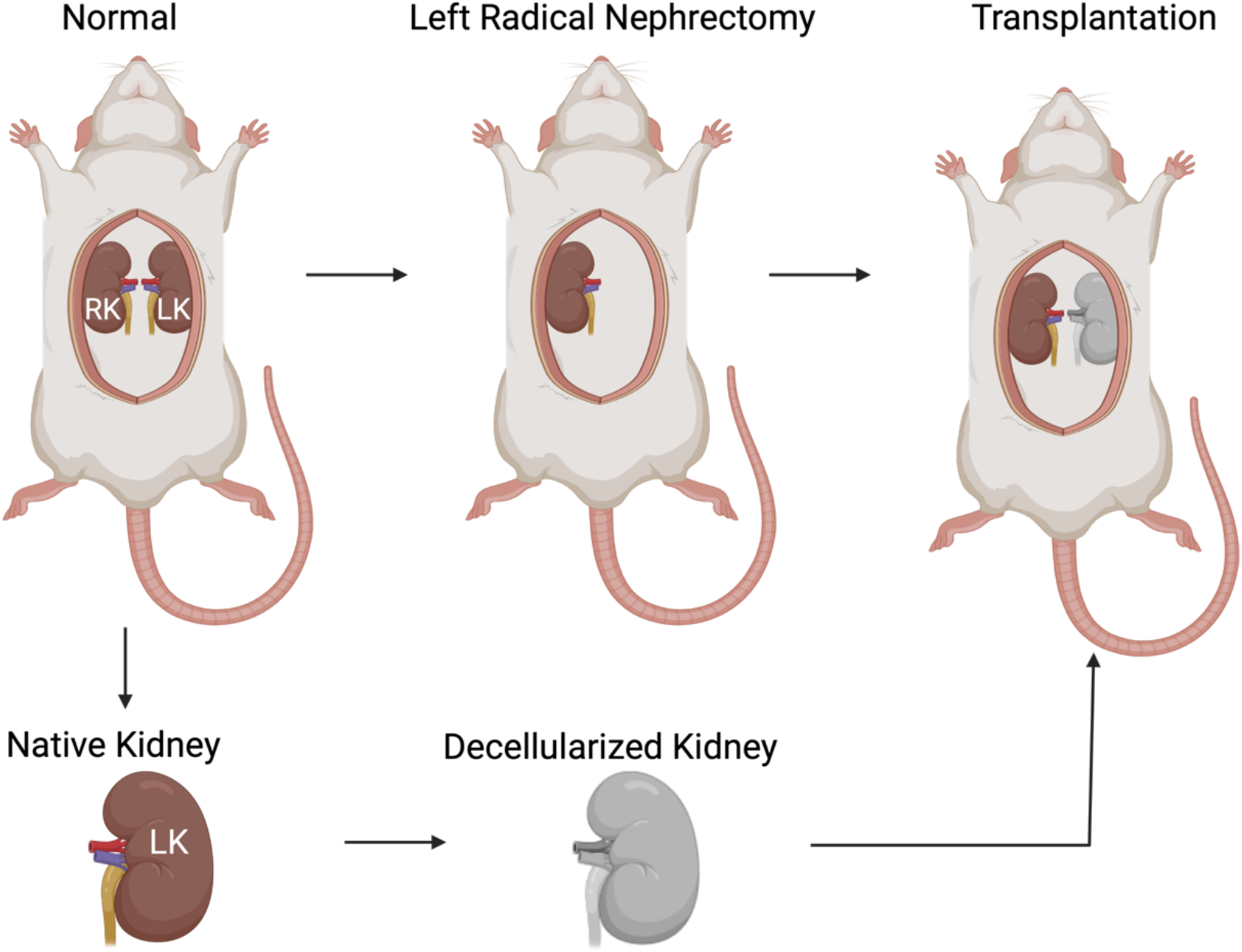
An illustration of the autologous and allogeneic orthotopic transplantation schema. Left-radical nephrectomies were first conducted to obtain kidneys with intact segments of renal arteries, veins, and ureters. Each native kidney was then decellularized and transplanted back into its respective recipient or implanted into another uninephrectomized recipient.

### 2.8 Intravital Two-Photon Microscopic Assessment of Nuclear Labeling, Capillary Blood Flow, Vascular Permeability

Exteriorized native or acellular kidneys were individually positioned inside a 50 mm glass-bottom dish (Willco Wells B.V., Amsterdam, The Netherlands) containing saline, set above an X60 water-immersion objective for imaging, and a heating pad was placed over the animal to regulate body temperature, Figure 2[40, 50]. The study was conducted using an Olympus FV1000MP multiphoton/intravital microscope (Center Valley, PA, USA) equipped with a Spectra-Physics (Santa Clara, CA, USA) MaiTai Deep See laser. The laser was tuned to excite Hoechst and FITC dyes across 770-860 nm excitation wavelengths. Images were collected with external detectors that acquired blue, green, and red emissions. The infusates were introduced via the tail vein. A schematic illustrating the intravital imaging process is presented in Figure 2.

**Figure 2.**
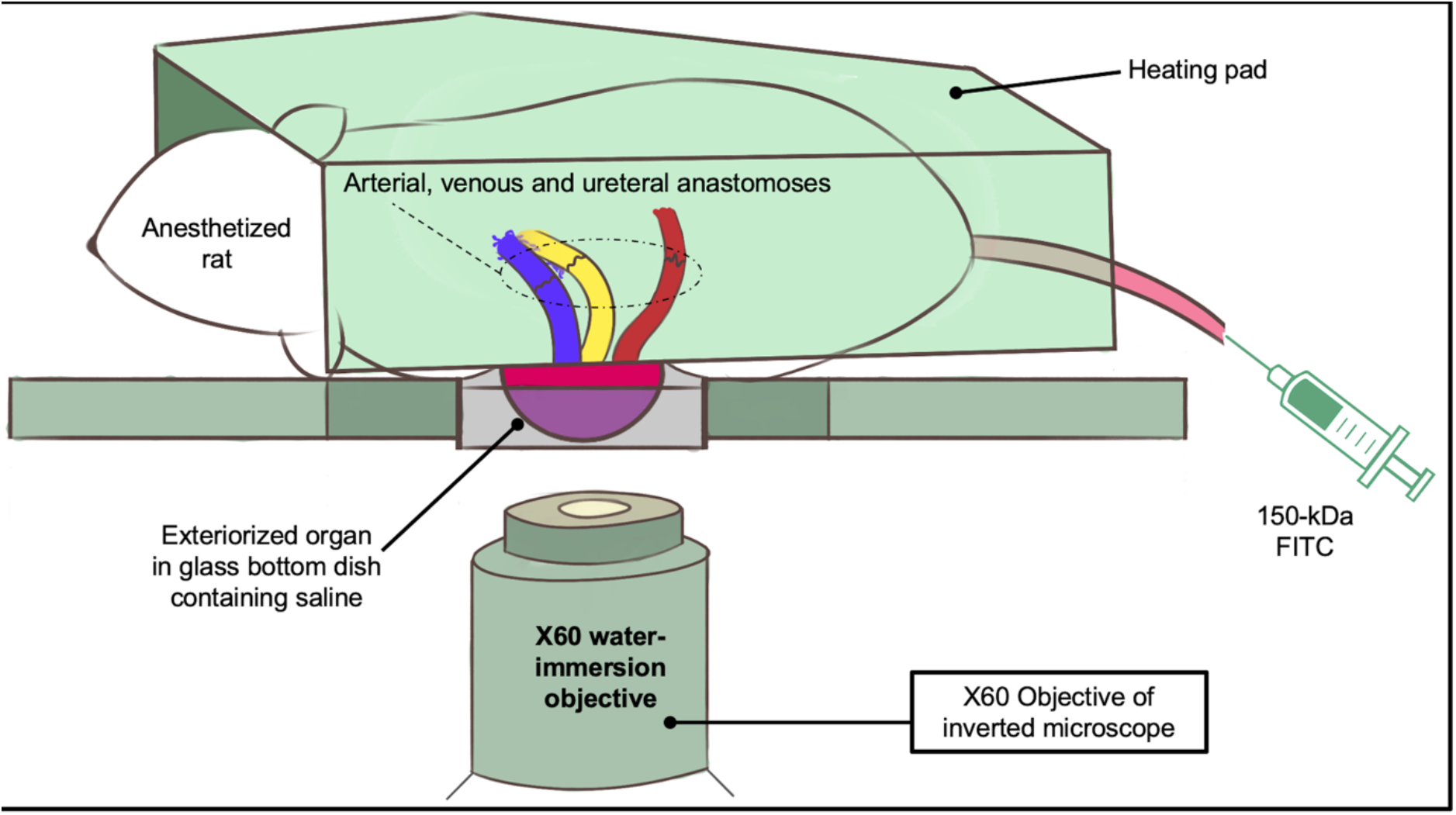
A schematic illustrating the intravital imaging process used to visualize live transplanted or native kidneys. As outlined in the literature, anesthetized rats with exteriorized (native or transplanted) kidneys were placed 50 mm glass-bottom dish, filled with saline, and set above the stage of an inverted microscope with a Nikon ×60 1.2-NA water-immersion objective. A 25-gauge butterfly needle was inserted into the dilated vein tail vein and attached to a syringe containing injectates. The heating pad was then placed directly over the animal to maintain core temperature.

Images were collected to analyze nuclear remnants in scaffolds, changes in tissue fluorescence, and microvascular integrity and function, as outlined in the literature[38, 39, 46, 47, 56], at 0-, 12-, 24- and 1-week time points. ImageJ software (Fiji- ImageJ × 64, US National Institutes of Health, Bethesda, MD, USA) was used to examine changes in nuclear DNA content, microvasculature permeability, and microarchitectural dimensions after transplantation by selecting four regions in various renal compartments at random to measure changes in the average fluorescence intensities at the defined measurement points, as previously outlined in the literature[39, 56]. The rouleaux density was determined from the number of stacked red blood cells in vascular compartments within adjacent fields divided by the vascular area, and glomerular diameters were also estimated using the ImageJ software.

### 2.9 Statistical Analysis

SPSS (IBM Corp, Armonk, NY, USA) was used to perform non-parametric analyses. The Kruskal-Wallis one-way analysis of variance with the post hoc Dunn’s test examined remnant DNA and SDS concentrations within acellular kidneys using biochemical or microscopic studies and differences in fluorescence intensities in various renal compartments between native and decellularized kidneys. This non-parametric test was also used to examine whether the degrees of dextran extravasation from the microvasculature and alterations to capillary blood flow directly after transplantation and at 12-hour, 24-hour, and 1-week time points were significant. All variables are expressed as mean ± standard deviation, and a p-value of less than 0.05 was considered statistically significant for all evaluations.

## 3 Results

### 3.1 Biochemical Assays Used to Evaluate the Decellularization Process

Substantial visible changes to the original organ (Figure 3A) occurred as early as 4 hours (Figure 3B) during decellularization, as it ultimately transitioned to a translucent structure (Figure 3C). The resulting scaffolds were then perfused with PBS to remove the cellular and detergent remnants (Figure 3D) and subjected to biochemical assays to evaluate residual DNA (Figure 3E) and SDS (Figure 3F) concentrations. The assays revealed that this method removed approximately 98% of the innate DNA content from the original kidneys and roughly 99% of the remnant SDS from scaffolds. The significant removal of DNA (Kruskal-Wallis, H = 23.298, d.f. = 3, p < 0.001) and SDS (Kruskal-Wallis, H = 23.345, d.f. = 3, p < 0.001) confirmed the effectiveness of this protocol.

**Figure 3.**
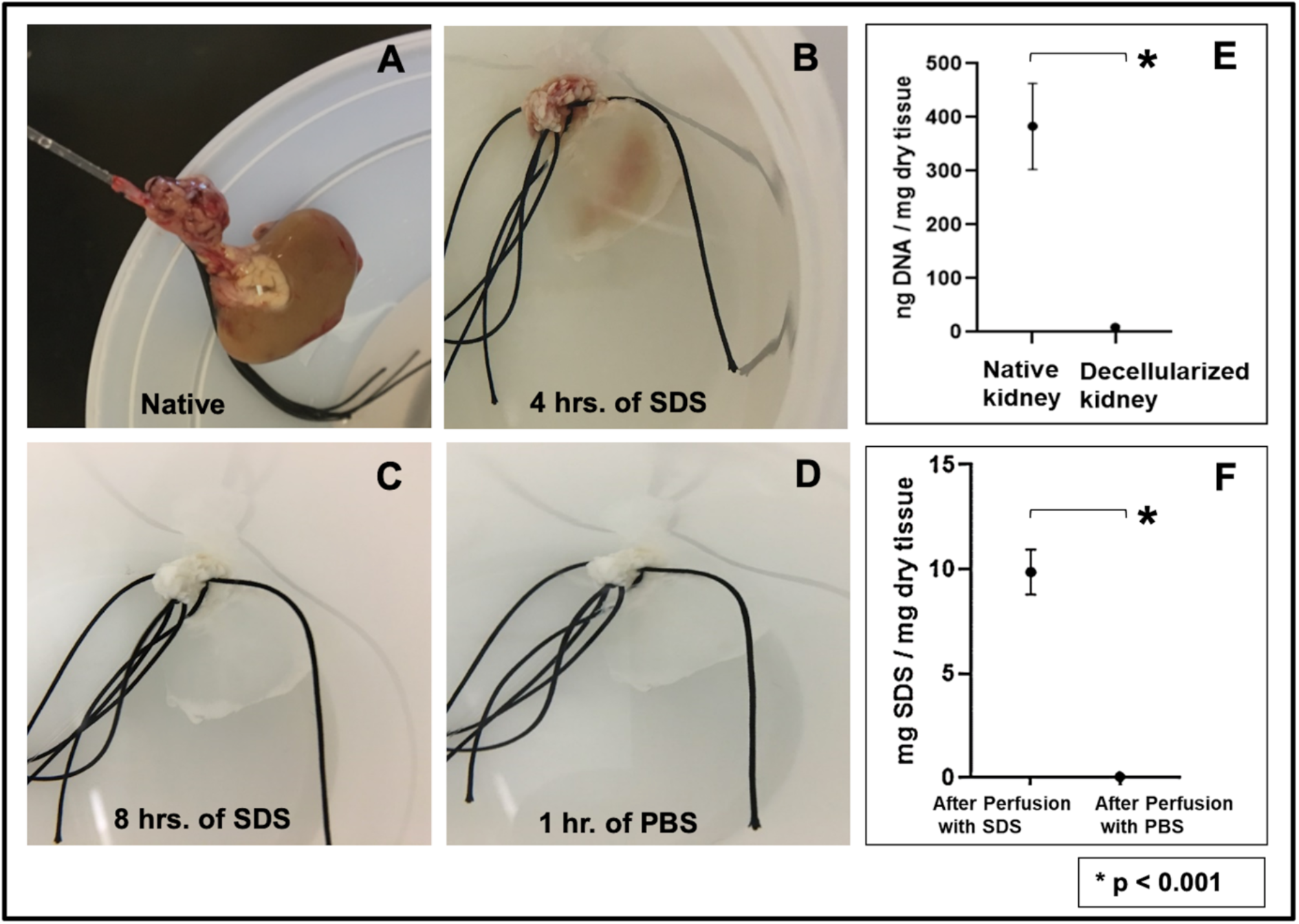
An evaluation of the decellularization process using biochemical assays. A) An image of a whole rat kidney with its renal artery cannulated for perfusion-based decellularization. B) The kidney, after 4 hours of perfusion with SDS, illustrates its transition from a solid to a translucent structure. C) The fully decellularized kidney is shown after 8 hours of SDS perfusion. D) The kidney after its subsequent perfusion with PBS. E) A plot presenting the low level of remnant DNA in the scaffold after decellularization. F) A graph highlighting the effective removal of SDS from the scaffold and its low residual concentration. Non-parametric evaluations conducted using Kruskal-Wallis detected significant declines in DNA and SDS concentrations after decellularization (* p < 0.001).

### 3.2 Whole Rat Kidney Decellularization Evaluated by Intravital Two-Photon Fluorescence Microscopy

Two-photon intravital micrographs, obtained from the blue pseudo-color channel, revealed considerable variations in fluorescence from native (non-transplanted) kidneys (Figure 4A) and decellularized scaffolds that were transplanted into live recipients after introducing Hoechst 33342 (Figure 4B). Measurements obtained from these scaffolds showed an approximate 92% drop in the relative level of blue-pseudo fluorescence, signifying the absence of nuclear staining compared to the native kidney. This significant reduction in the relative blue pseudo-color fluorescence (Kruskal-Wallis, H = 25.114, d.f. = 3, p < 0.001) correlated with the decrease in DNA content obtained from the biochemical assay (Figure 3E).

**Figure 4.**
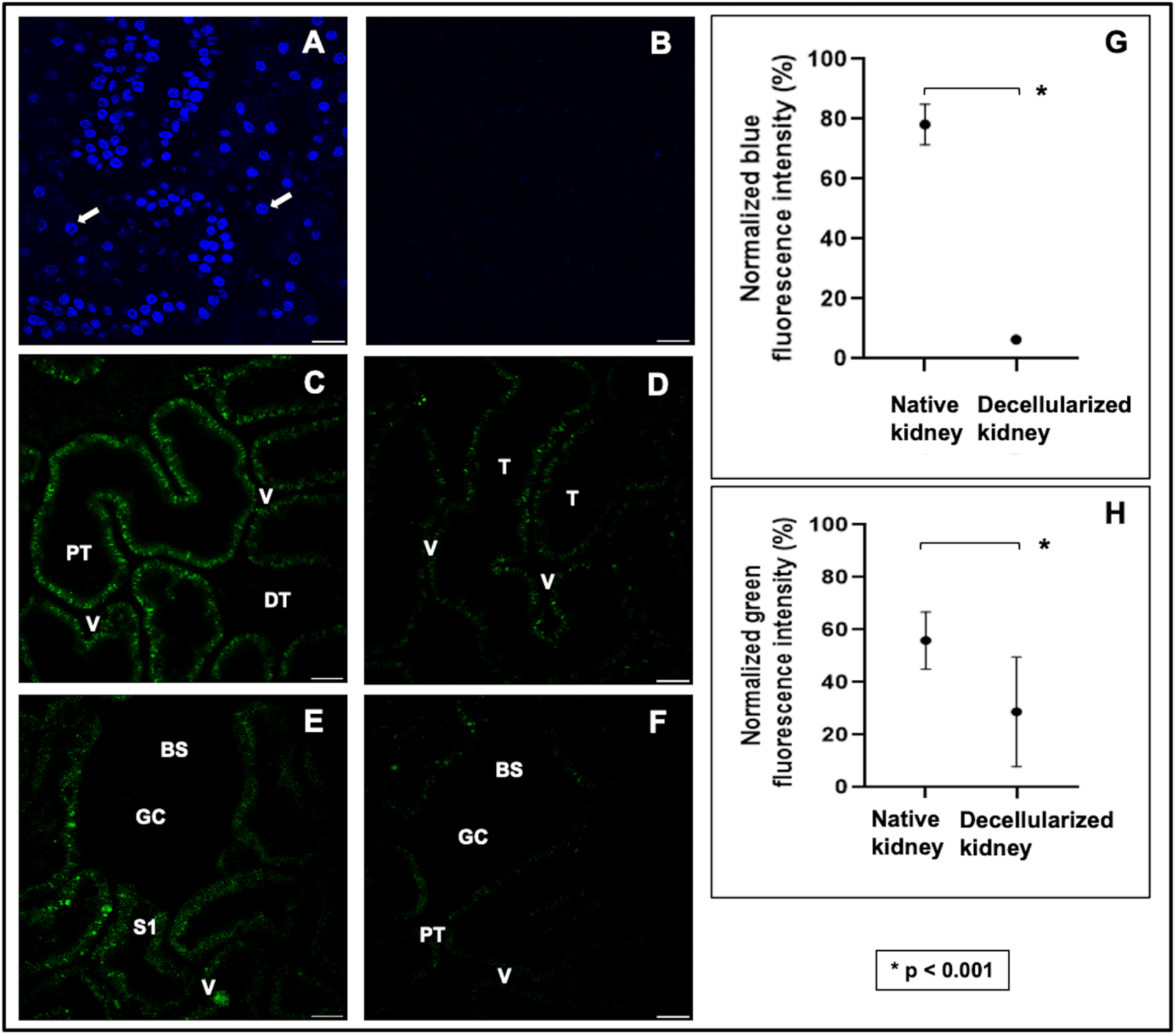
An evaluation of the decellularization process using intravital microscopy. A) Images obtained from a native (non-transplanted) kidney show the nuclear stain’s vibrant presence. B) Image from a transplanted decellularized kidney showing the absence of Hoechst 33342 labeling and comparing green autofluorescence signals gathered from native (non-transplanted) and transplanted decellularized kidneys. C) Image obtained from a native kidney that presents only distal (DT) and proximal (PT) tubular compartments and peritubular vascular tracks (V). D) Intravital micrograph identifying decellularized tubular (T) and vascular compartments. It should be noted that the decellularization process made it difficult to differentiate between tubular segments. E) Image obtained from a native kidney that captured the Bowman’s space and glomerular capillaries (GC), the S1 segment of the proximal tubule compartment (S1), and peritubular vasculature. F) Image of a scaffold kidney highlighted the decellularized glomerular and tubular segments. G) Comparison of relative blue pseudo-color fluorescence intensity from native and decellularized scaffolds. H) Comparison of relative green pseudo-color fluorescence intensity from native and decellularized scaffolds. Non-parametric evaluations conducted using the Kruskal-Wallis test detected significant reductions in both the normalized blue (* p < 0.001) and green (** p < 0.001) pseudo-color fluorescence observed after decellularization. Scale bars represent 20 μm.

Similarly, images collected from the green pseudo-color channel displayed variations in the intrinsic level of autofluorescence present in native kidneys (Figure 4C and Figure 4E) compared to that of the decellularized organ (Figure 4D and Figure 4F). Within these structures, proximal tubules were quickly identified as having the most outstanding autofluorescent signal and thickness, whereas distal tubules appeared thinner and dimmer (Figure 4C). Likewise, the characteristic shape of the renal corpuscle was highlighted by the outline of the Bowman’s capsule and faint or invisible capillary tuft, along with the nearby peritubular capillary and interstitial space (Figure 4E). However, after decellularization, significant (Kruskal-Wallis, H = 24.353, d.f. = 3, p < 0.001) reductions in both the blue (Figure 4G) and green (Figure 4H) pseudo-color signal intensities were observed. These drops in the relative fluorescence levels in the scaffolds made it challenging to identify specific tubular compartments (Figure 4D), except proximal tubule segments that emanated from the Bowman’s capsule (Figure 4F).

### 3.3 Real-time In Vivo Examination Show Alterations to Blood Flow and Dextran Extravasation within Scaffolds Directly after Transplantation

In vivo data showed that the fluorescence levels in the tubular epithelial and luminal compartments remained relatively constant and comparable in both native (Figures 5A-5C and 5G) and decellularized (Figures 5D-5F and 5H) kidneys within the first few minutes after transplantation. Specifically, fluorescence intensity levels within the vascular lumens of native and decellularized kidneys were substantially higher than the intensities in the other mentioned renal compartments post-transplantation. The Kruskal-Wallis test revealed that these differences within native kidneys were significant (Kruskal-Wallis, H = 36.028, d.f. = 1, p < 0.001), and the post hoc Dunn’s test only detected significant pairwise differences between the following: fluorescence intensity in the vascular lumen and the tubular epithelium (Kruskal-Wallis, H = 36.028, d.f. = 1, p < 0.001), and vascular lumen and the tubular lumen (Kruskal-Wallis, H = 34.392, d.f. = 1, p < 0.001).

**Figure 5.**
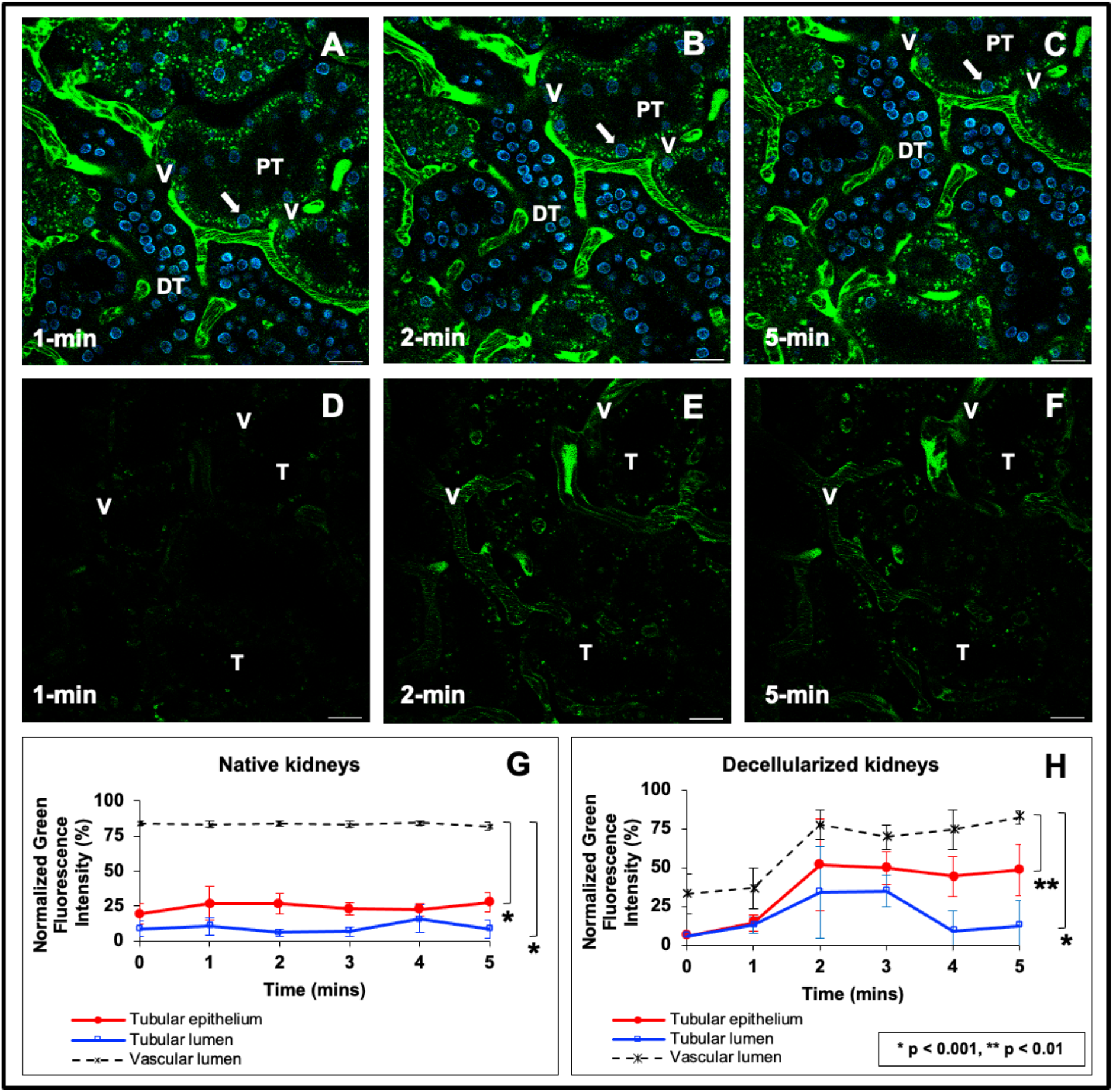
In vivo assessment of microvascular leakage in decellularized nephrons directly after transplantation. A) through C) Time-series images taken across 5 minutes from a live native kidney display the proper confinement of large molecular weight dextrans within the peritubular vasculature (V) and characteristic autofluorescence levels within proximal and distal tubules and nuclear staining with Hoechst 33422 (arrows). D) through F) Images obtained at 1 min, 2 min, and 5 min marks from a transplanted acellular scaffold also display the confinement of large molecular weight dextrans within the peritubular vasculature (V) and substantially reduced autofluorescence levels within the tubules, and absence of nuclear staining. Tubular lumens are highlighted by (L). Scale bars represent 20 μm. G) A graphical representation of green pseudo-color fluorescence variations outlining relatively constant and elevated fluorescence levels within the vasculature, compared to the much lower signals recorded in the tubular lumen and epithelium. H) A similar graphical representation of the variations in green pseudo-color fluorescence levels observed in decellularized kidneys shows the rising level and FITC fluorescent within the vasculature compared to lower signals recorded in the tubular lumen and epithelium. The data presented in this figure highlight events recorded from autologous transplantation and are comparable to those observed in allogeneic cases.

Likewise, for decellularized kidneys, this non-parametric test also detected a significant difference in the intensities in the tubular epithelia, tubular lumen, and vascular lumen (Kruskal-Wallis, H = 12.487, d.f. = 1, p < 0.001). Thereafter, the ad hoc test only detected significant pairwise differences between the fluorescence intensity in the vascular lumen and the tubular epithelium (p < 0.01), vascular lumen and the tubular lumen (Kruskal-Wallis, H = 26.508, d.f. = 1, p < 0.001), and tubular epithelium and the tubular lumen (Kruskal-Wallis, H = 7.362, d.f. = 1, p < 0.007). The dextran molecules were exclusively present in vascular lumens immediately after scaffold transplantation. However, there was a delayed, reduced, and inhomogeneous distribution of the dextrans in the decellularized vasculature (Figures 5D-5F and 5H) compared to the distribution in normal kidneys imaged under the same conditions (Figures 5A-5C and 5G). It is conceivable that additional time was needed to fill the acellular nephrons with blood directly after transplantation. Our statistical analyses also detected that the variations between blood quantities within native and decellularized peritubular capillary tracks significantly differed within this period (Kruskal-Wallis, H = 13.505, d.f. = 1, p < 0.001). These quantities of blood were indirectly estimated by the dextran fluorescence intensity within the microvascular lumen and highlighted signs of scaffold leakage. This observation can support the possibility of vascular leakage within the short term, which could have supported dye translocation that increased fluorescence levels in the decellularized epithelial and luminal segments. Moreover, no significant variations were detected in distinguished events related to autologous and allogeneic transplantation.

### 3.4 Severe Dextran Extravasation and Alterations to Capillary Blood Flow Observed Across One Week of Transplantation

In comparison to the immediate events after transplantation, drastic changes in vascular structure and function from the initial state (Figure 6A) were detected at the 12-hour (Figure 6B), 24-hour (Figure 6C), and 1-week (Figure 6D) time points. The varying degrees of dextran extravasation from the microvasculature observed across this period signified the impairment of typical filtrative capacities and increased permeability of the decellularized microvasculature in vivo. Correspondingly, from a quantitative perspective, measurements from these micrographs highlight one and two orders of magnitude increases in the presence of the dye within the Bowman’s space (Figure 7A), tubular epithelium (Figure 7B), tubular lumen (Figure 7C), interstitial space (Figure 7D), and peritubular capillary endothelium (Figure 7E). Alterations to capillary blood flow outlined by the increased presence of rouleaux were also investigated post-transplantation. Rouleaux density measurements indicated one and two orders of magnitude increases in red blood cell aggregation (Figure 7F). Finally, estimations of glomerular diameter (Figure 7G) provided evidence of substantial increases across the measurement period. Overall, these analyses revealed statistically significant pairwise differences in the quantities of the vascular marker extruded from the capillaries and the changes in rouleau density and glomerular diameter across the 7-day measurement period, again without being able to identify significant differences between the autologous or allogeneic nature of the transplantation.

**Figure 6.**
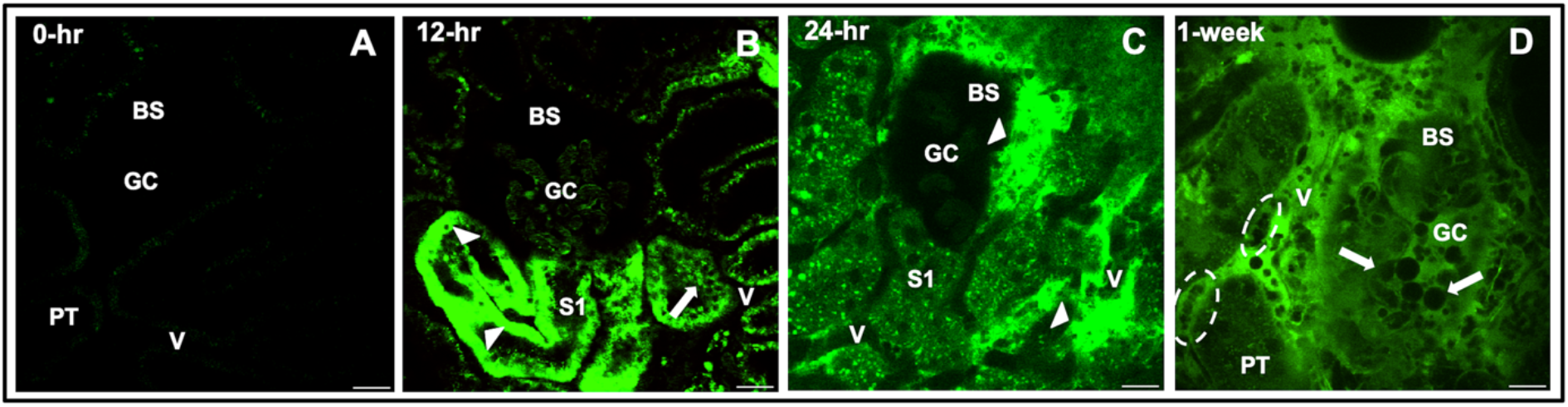
Disruptions to scaffold integrity during a week after transplantation. A) Image taken from a decellularized kidney that displays the decellularized autofluorescence before the introduction of FITC (this is the same imaging field that is shown in Figure 4F). B) Image taken from a decellularized kidney 12 hours after transplantation illustrates substantial and inhomogeneous levels of dye translocation between luminal, epithelial, and interstitial compartments. C) Image taken from a decellularized kidney 24 hours after transplantation presents the accumulation of the dextran and blebs within the Bowman’s capsule, interstitium, and tubules. D) Image taken from a transplanted decellularized kidney 1 week (168 hours) after transplantation provides evidence of rouleaux (dashed oval within the vasculature) and bleb/vesicle (arrows within the Bowman’s space and tubular lumen) formation that accompanied dye extrusion from breached decellularized glomerular capillary tracks to completely occlude this enlarged glomerulus. Scale bars represent 20 μm. The data presented in this figure highlight events recorded from autologous transplantation and are comparable to those observed in allogeneic cases.

**Figure 7.**
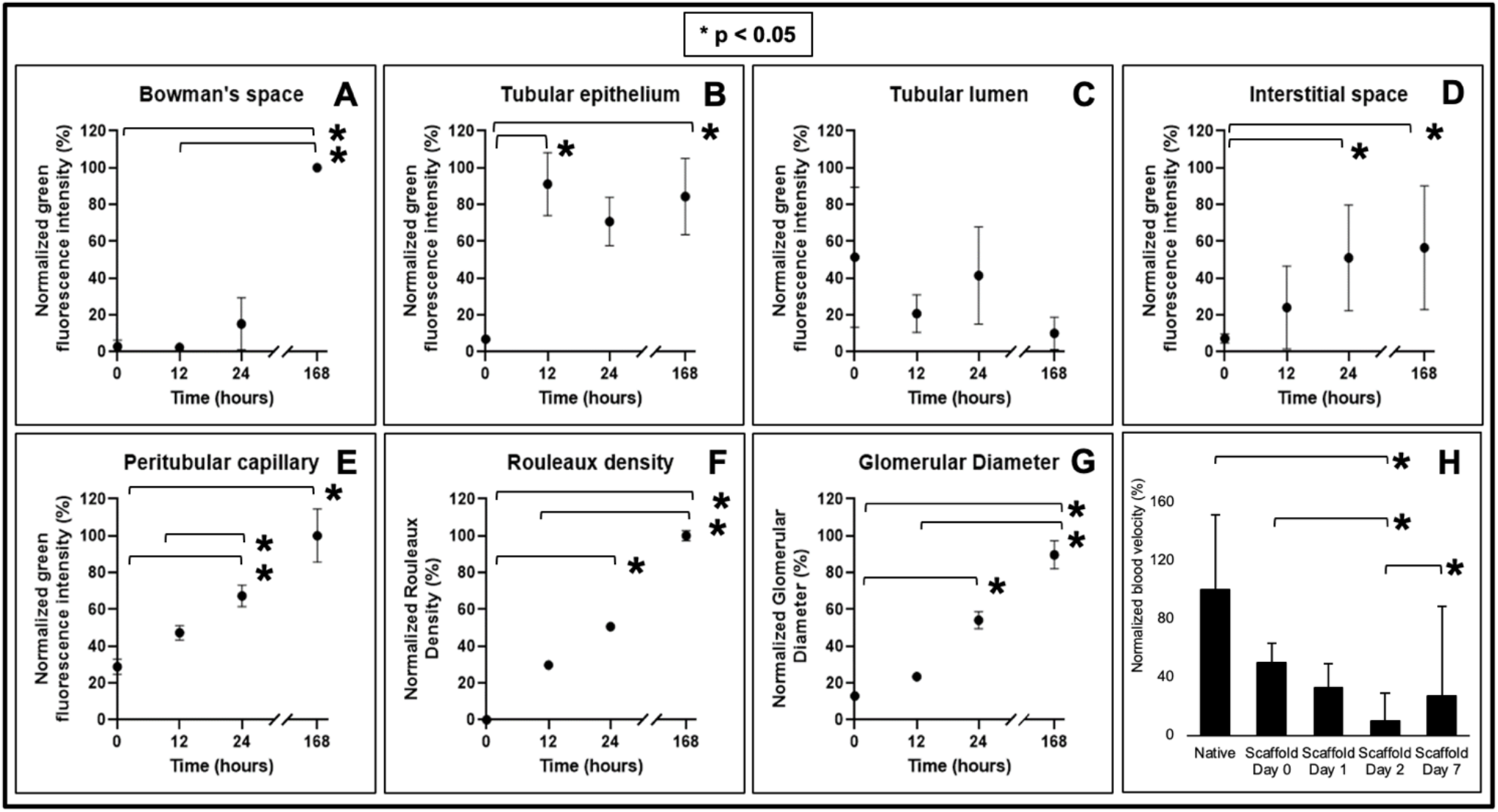
Estimations of blood extravasation from various decellularized renal compartments, rouleaux density, glomerular hypertrophy, and velocity within the microvasculature. Graphs examining the degree of FITC dye translocated within A) the Bowman’s space, B) tubular epithelium, C) tubular lumen, D) interstitial space, E) peritubular capillary endothelium, as well as F) rouleaux density and G) glomerular diameter, during the 168-hour measurement period. H) In vivo assessment of velocity within the microcirculation of native kidneys and transplanted acellular scaffolds.

### 3.5 In vivo assessment of velocity within the microcirculation of native kidneys and transplanted acellular scaffolds

The velocities within the microcirculation of native kidneys and acellular scaffolds at day 0 (0 hrs.), day 1 (24 hrs.), day 2 (48 hrs.), and day 7 (168 hrs.) were estimated. The velocities were computed as ratios of blood displacement and time and expressed as relative values based on the rates observed within the native microvasculature. This assessment revealed substantial reductions in the velocities within the microcirculation observed within the scaffold. Analyses from the one-way non-parametric ANOVA detected a significant difference among velocities within microcirculatory systems in the abovementioned cases (p < 0.019), and Dunn’s post hoc test revealed pairwise differences between the velocity of the microcirculation of native kidneys and scaffolds on day 2 (p = 0.002), scaffolds on day 0 and scaffolds on day 2 (p = 0.012), scaffolds on day 2 and scaffolds on day 7 (p = 0.019). These results can be found in Figure 7H.

## 4 Discussion

Several approaches to developing a bioartificial kidney have been demonstrated, and whole organ decellularization appears to be the most promising methodology thus far[57]. One major challenge to this strategy is maintaining vascular integrity and functionality post-transplantation. Specifically, There needs to be a better understanding of how the integral components of the scaffold can be used to create complex bioartificial organs which can withstand post-transplantation environments. Developing solutions to this problem will identify ways to promote scaffold longevity and angiogenesis in bioartificial organs. A study from our group previously illustrated the effects generated by autologous transplantation of decellularized scaffolds. This study extends to the original and complementary work by examining events related to autologous and allogeneic transplantation. Such an extension is valuable and insightful because, in practice, organ transplantation occurs between different individuals. Thus, parameters limiting scaffold implantation in both instances have been identified. Most models used to examine the microvasculature have primarily utilized in vitro or in vivo techniques incapable of providing adequate spatial and temporal resolution. Thus, IVM has been employed to help unravel other vital mechanisms to understand this complex issue better. This advanced imaging technique aids the monitoring of live physiological/pathophysiological cellular and subcellular events in real-time[58], which can provide a better understanding of the events that adversely alter scaffolds post-transplantation.

Whole kidneys were extracted from rats and decellularized by perfusion, and the scaffolds were then transplanted into their respective donors. After that, nuclear and vascular dyes provided real-time evidence of the effective removal of cell/nuclear remnants from the acellular scaffolds and their structural and functional integrities. Evaluating the concentrations of these residual components is a crucial aspect of the decellularization process. It is necessary to efficiently remove cellular remnants from scaffolds to reduce the potential for unwanted immunological responses or tissue rejection after transplantation[59]. It is also essential that residual concentrations of SDS be low enough so that the detergent does not continue to denature the scaffold significantly. This unwanted effect can alter the scaffold’s overall permeability and compromise its integrity, adversely affecting recellularization and transplantation[60].

Intravital two-photon microscopy was used to examine the effectiveness of the decellularization process in vivo. Blue pseudo-color channel multiphoton micrographs revealed considerable variations in Hoechst 33342-based fluorescence between native and decellularized kidneys. This cell membrane-permeant dye diffuses readily into tissues[39-42, 54, 56]. It labels DNA in live and fixed cells by binding to adenine-thymine-rich DNA regions in the minor groove to produce extensive enhancements in fluorescence[61]. Such enhancements support the clear visualization of nuclei in the tubular endothelia and epithelia, glomeruli, interstitial cells, and circulating leukocytes[40]. Interestingly, similar drastic changes in tissue fluorescence were previously observed using immunohistochemical techniques in vitro and confirmed decellularization[22, 60, 62]. Thus, these in vivo results complement conventional in vitro histological and fluorescent microscopic methods routinely applied to confirm decellularization[63], highlighting the effective decellularization outlined by Crapo et al[28].

Similarly, images collected from the green pseudo-color channel displayed variations in the intrinsic levels of autofluorescence present in native kidneys. Typically, the kidney has a high level of green autofluorescence[39, 56], characterized by the natural emission of light by biological structures such as mitochondria and lysosomes[49], and unique metabolites like aromatic amino acids, nicotinamide adenine dinucleotide (NADH and its phosphate analog NADPH), and flavins[47, 64]. These structures’ relative distribution and percentages vary and help distinguish renal compartments, namely the proximal and distal tubules, glomerulus, peritubular capillaries, and interstitium, without the addition of fluorescent markers[19, 40].

Once decellularization was established, IVM was utilized to examine the structural and functional integrities of the scaffolds post-transplantation. Scaffold integrity was first investigated immediately post-transplantation, whereby time-series data was used to monitor the live introduction of the 150-kDa FITC dextran in the decellularized nephron. This dextran is a branched polysaccharide. The fluorescein moiety is attached by a stable thiocarbamoyl linkage, which does not lead to any depolymerization of the polysaccharide and helps the dextran retain its minimal charge. These features ensure that the fluorescent molecules are restricted to the vasculature and are essential for permeability studies.

Overall, the data provided real-time evidence that these large molecular weight molecules were primarily confined to the decellularized vascular lumen directly after transplantation. However, the data also suggested that the dye began to leak through the modified renal filtration and peritubular barriers (Figure 5D through Figure 5F) compared to native, non-transplanted kidneys (Figure 5A through Figure 5C). This view is based on the relative variations in fluorescence levels observed in the tubular epithelial of the native and decellularized kidneys during this period previously stated, as well as variations between signals observed in the native and decellularized lumens. Such results emphasize the power of intravital microscopy to uncover subtle changes in vascular permeability, even with a small sample size. Moreover, the explanted decellularized kidney vasculature could initially withstand some degree of in vivo blood flow/pressure levels, and lower degrees of dye translocation was detected at the microscopic level.

Vascular permeability was further investigated throughout the week that followed transplantation. For this investigation, the 150-kDa FITC dextran was infused into the tail vein directly before blood was introduced into transplanted scaffolds. This process provided a means to evaluate the distribution of the blood constituents and insight into the in vivo environment’s effect on the grafts. This infusion regimen was favored over infusing the dye at later times. It was necessary to ensure that the fluorescent marker circulated through the decellularized nephron because, at future time points, clotting could prevent the dye from entering the decellularized microvasculature.

Ordinarily, the native kidney can autoregulate blood flow to safeguard against considerable fluctuations in blood pressure transmitted to peritubular and glomerular capillaries[65]. However, the acellular organ cannot replicate this vital function[19]. As a result, scaffolds would have been exposed to abnormal pressures capable of damaging these more delicate decellularized structures[7]. Mechanistically, such damage could explain the progressive leakage of the FITC dye from the glomerular capillaries into the Bowman’s space (Figure 8). Decellularization would have actively altered microarchitectural permeability, disrupting the decellularized glomerulus’ ability to act as a natural sieve that limits the passage of only water and small solutes into the filtration pathway.

**Figure 8.**
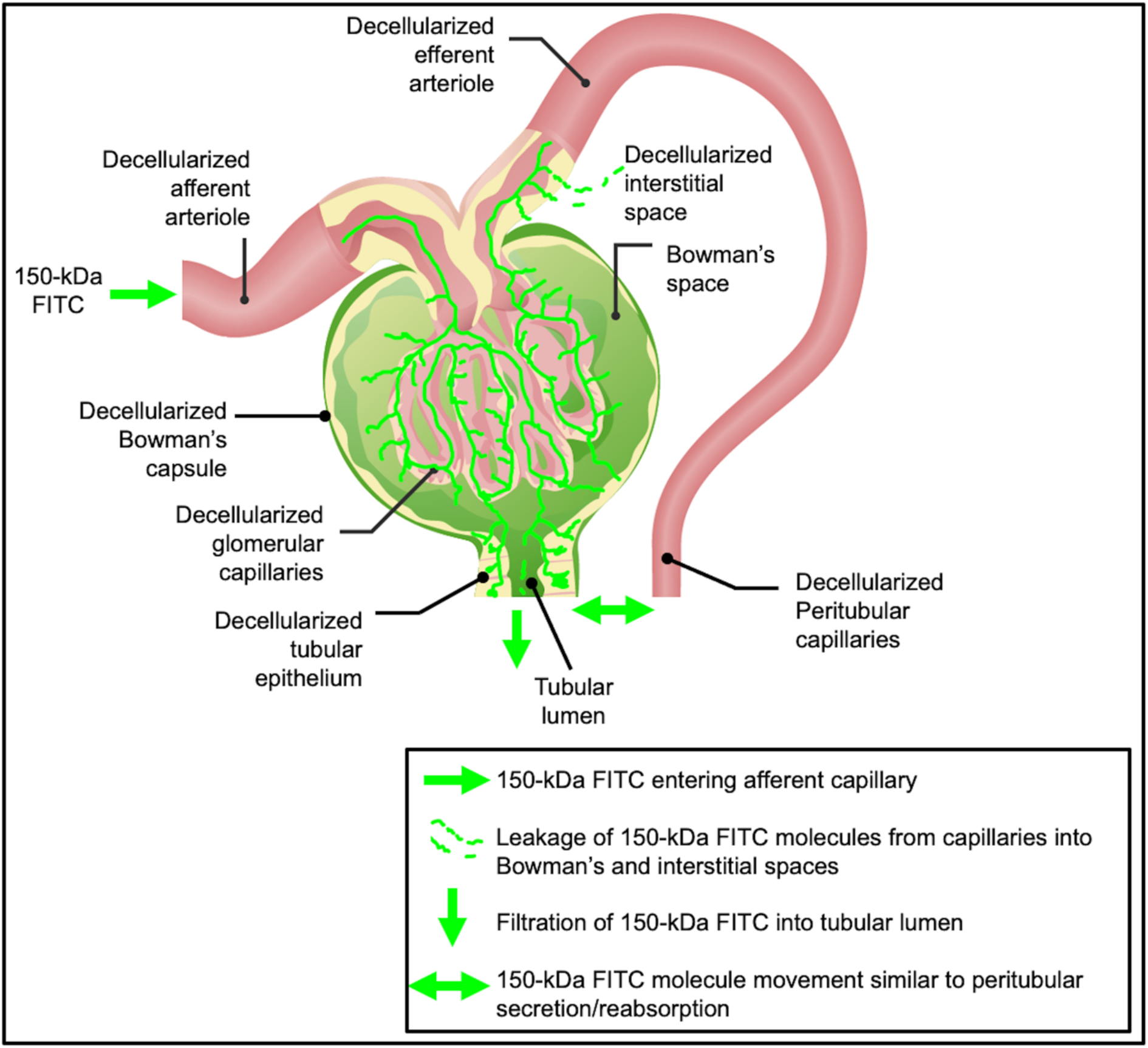
A proposed simplified view of dextran extravasation from the decellularized vasculature and translocation to the extravascular space. This model presents the large molecular weight, 150-kDa FITC, dye’s entry into the decellularized nephron vasculature via the afferent arteriole. The vascular marker then progresses through the decellularized glomerulus, where it can be filtered into the Bowman’s space and enter the tubular lumen. This unregulated process can potentially induce significant dynamic and static pressures to facilitate bilateral dye translocation between tubular epithelium, interstitium, and peritubular endothelium. The original image was adapted with permission[69].

Additionally, within the native glomerulus, the fenestrated capillary endothelial barrier restricts the passage of molecules smaller than 70 kDa[66]; the basement membrane, with its negative charge, also limits the passage of particles movement and favors the filtration of cations. Foot processes on podocytes provide an additional size selectivity as they wrap around the glomerular capillary loop to form interdigitated filtration slits with a spacing of approximately 40 nm[67]. By removing this intrinsic selectively permeable barrier through decellularization, macromolecular transport was no longer limited by size, shape, charge, and deformability[68].

With the loss of the characteristics mentioned above, the acellular glomerulus would likely have facilitated the filtration of the 150-kDa molecules into the Bowman’s space and tubular lumen. The accumulation of blood within these regions would have created opportunities for blood to extravasate into the denatured acellular tubular epithelium and interstitium. Thus, such fluid translocation would likely have mimicked unregulated secretion and reabsorption patterns throughout the fractured nephron. Specifically, this dye would have subsequently entered the lumens of the decellularized peritubular capillaries and tubular epithelium after filtration. Furthermore, the simultaneous accumulation of fluid, macromolecules, and blood cells within these compartments could have established uncharacteristic hydrostatic, hydrodynamic, and osmotic pressure gradients. These pressures could have, in turn, supported the translocation into epithelial/interstitial compartments shown in the intravital micrographs (Figure 5B through Figure 5D). This effect could have undoubtedly reduced the dye concentration within the patent regions of the vasculature, as evidenced within glomerular capillaries at the 12-hour and 1-week marks.

Apart from this, erythrocytic aggregation is a hallmark of ischemia[38]. This outcome was anticipated, as collagen within transplanted scaffolds would have been exposed to flowing blood. Such an interaction would have rapidly facilitated platelet activation and aggregation[70]. Within the ECM, collagen is the only protein that supports both platelet adhesion and complete activation[70]. Collagens within this matrix are generally separated from blood by the endothelial layer. However, the decellularized scaffold directly contacts flowing blood and effectively initiates hemostasis. It is expected that these processes were unregulated in the transplanted scaffolds and generated widespread coagulation and thrombosis, which would have ultimately facilitated the reductions in blood flow and entrapment of the dye within acellular nephrons observed at the 1-week mark.

These mechanisms are also known to support bleb and microvesicle formation visualized in vivo within decellularized glomerular segments. Bleb/microvesicle accumulation within the microvasculature can lead to longstanding occlusions, particularly in decellularized vessels that cannot compensate for fluctuations in blood pressure. Furthermore, unresolved obstructions to blood flow generate detrimental stagnation pressures that can induce antegrade blood flow and renal overload compartments like the glomerulus. This process can lead to decellularized glomerular hypertrophy, outlined by marked increases in the size of the decellularized renal corpuscle (Figure 6D) compared to that of the native (Figure 6A). This form of hypertrophy could have arisen from considerable changes in efferent and afferent flows/pressures exerted on acellular glomeruli[71]. Excessive filtration and proteinuria could have also contributed to the condition[72]. Cellular debris and vascular cast formation in the Bowman’s space that could have arisen from ischemia-derived blood cell apoptosis or necrosis observed in chronic diseases are also likely contributing factors[73]. Signs of these effects were observed by tracking the changes in glomerular diameter, which markedly increased with time (Figure 7G).

Finally, it is also essential to recognize that intravital microscopy studies on the glomerulus are typically conducted in Munich-Wistar rats with superficial glomeruli, which can be routinely accessed by intravital two-photon microscopy[47, 51]. Surface cortical glomeruli are scarce in Sprague-Dawley rats[45], and intrinsic tissue autofluorescence generally inhibits live imaging in the kidney beyond 150 μm[43]. Interestingly, the decellularization process substantially reduced the tissue autofluorescence and provided access to multiple glomeruli throughout the study, emphasizing the acquired ability to image deeper in the acellular organ and visualize these previously hidden structures in the Sprague-Dawley native kidney. Altogether, these results indicate that an in vivo method capable of tracking microvascular integrity represents a powerful approach for studying scaffold viability and identifying ways to promote scaffold longevity and angiogenesis in bioartificial organs.

## 5 Conclusion

Even though the optimal conditions to achieve a decellularized whole organ have yet to be devised, current best practices have outlined SDS as a primary agent for creating acellular scaffolds. Using this single agent, we effectively decellularized rat kidneys and highlighted the use of intravital fluorescent microscopy to assess scaffold generation and structural and functional integrity.

Specifically, these studies showed that scaffolds orthotopically transplanted into rats initially retained a reasonable degree of microvascular structure in vivo directly after transplantation. However, the scaffold then succumbed to widespread coagulation and thrombosis, which would have eventually facilitated the reductions in blood flow within acellular nephrons. Simultaneously, the loss of intrinsic barriers and compensatory mechanisms provided additional means to further hamper in vivo viability. Altogether, valuable insight into blood filtration and extravasation mechanisms was obtained and provided sites within the decellularized nephron that need to be structurally reinforced and a possible timeline during which devastating changes occur. This approach can also identify ways to maintain viable blood supply and limit scaffold degradation in post-implantation environments. Such an understanding will support the next stage in the evolution of bioartificial organs.

## 6 Data Availability Statement

The datasets generated during and/or analyzed during the current study are available from the corresponding author on reasonable requests.

## 7 Ethics Statement

The animal study was reviewed and approved by the Institutional Animal Care and Use Committee (IACUC) at the School of Medicine, Wake Forest University; Animal Research Oversight Committee (AROC) at Khalifa University of Science and Technology and aligned with ARRIVE criteria.

## 8 Author Contributions

P.R.C. conceived and designed research, performed experiments, analyzed data, interpreted results of experiments, and prepared the manuscript.

## 11 Acknowledgments

The author acknowledges funding from an Institutional Research and Academic Career Development Award (IRACDA), Grant Number: NIH/NIGMS K12-GM102773, and funds from Khalifa University, Grant Numbers: FSU-2020-25 and RC2-2018-022 (HEIC). The author would like to thank Dr. Joao Paulo Zambon, Dr. Amanda Dillard, and Mr. Kenneth Grant for their support in developing the decellularization process, transplant model, and imaging protocol. The author also wishes to thank Ms. Imaan Khan for creating the graphic presented in Figure 7 and Ms. Anousha Khan for technical help. Finally, the author thanks Mrs. Maja Corridon, Ms. Xinyu Wang, and Dr. Adeeba Shakeel for reviewing the manuscript.

